# Growing DAGs: Optimization Functions for Pathway Reconstruction Algorithms

**DOI:** 10.1101/2022.07.27.501737

**Authors:** Tunç Başar Köse, Jiarong Li, Anna Ritz

## Abstract

A major challenge in molecular systems biology is to understand how proteins work to transmit external signals to changes in gene expression. Computationally reconstructing these signaling pathways from protein interaction networks can help understand what is missing from existing pathway databases. We formulate a new pathway reconstruction problem, one that iteratively grows directed acyclic graphs (DAGs) from a set of starting proteins in a protein interaction network. We present an algorithm that provably returns the optimal DAGs for two different cost functions and evaluate the pathway reconstructions when applied to six diverse signaling pathways from the NetPath database. The optimal DAGs outperform an existing *k*-shortest paths method for pathway reconstruction and the new reconstructions are enriched for different biological processes. Growing DAGs is a promising step towards reconstructing pathways that provably optimize a specific cost function.

## 1 Motivation

Intracellular signaling pathways describe the molecules and interactions that convert a particular external signal (such as growth, proliferation, movement, or death) to the change of expression of one or more genes, culminating in a cellular response through transcriptional regulation. Many signaling pathway databases such as Reactome [1], KEGG [2], and NetPath [3] document the interactions associated with signaling pathways, and efforts such as WikiPathways [4] and Pathway Commons [5] have unified these databases by combining dozens of pathway resources. However, even pathways that describe fundamental biological processes or pathways implicated in commonly studied diseases are not complete: they are likely missing canonical proteins and protein interactions. Further, it is difficult to use these pathway databases to study less well-known signaling events. Computationally reconstructing a signaling pathway of interest from experimental data would be a huge help for the (mostly manual) curation of pathway databases such as Reactome and KEGG, and would provide a new lens to investigate signaling pathways that are not yet cataloged in these databases.

We and others have made use of large protein-protein interaction datasets as well as the annotated pathway databases to construct an *interactome* – a graph representation of physical interactions among all proteins in the organism under study. Since some interactions (especially those from existing databases) represent post-translational modifications where a direction of signal is clear, we consider an interactome as a weighted, directed graph *G* = (*V, E*) where the edges are weighted according to the supporting evidence.

### 1.1 Pathway Reconstruction Problem

In earlier work [6], we formulated the following **Pathway Reconstruction Problem**: Given a weighted, directed interactome *G* = (*V, E*), a set *R* ⊂ *V* of membrane-bound receptors specific to a pathway of interest, and a set *T* ⊂ *V* of transcriptional regulators specific to a pathway of interest, return a subgraph *G*′ ⊆ *G* that corresponds to the signaling pathway that connects nodes in *R* to nodes in *T*. We showed that a *k*-shortest paths (KSP) approach, PathLinker, outperformed existing methods for connecting nodes within a network to reconstruct pathways in pathway databases. There are two main reasons why a KSP approach such as PathLinker reconstructed pathways better than other methods. First, a KSP approach has a parameter to smoothly “grow” the network. Many existing algorithms return a small reconstruction because they aim to minimize the number of extraneous edges (such as Steiner forests [7] and network flow [8]). In KSP, we can increase the number of paths *k* to return iteratively larger reconstructions, capturing more of the pathway. Second, a KSP approach is guaranteed to connect nodes in *R* to nodes in *T*. While some methods such as Random Walks with Restarts (RWR) [9] have a parameter that “grows” the network, they are not guaranteed to connect the nodes from *R* to *T*. In a KSP approach, all paths start and end within the pathway by construction. Finally, reconstructions that are comprised of shortest paths are useful for generating hypotheses for follow-up validation. For example, we experimentally validated a path (RYK-CFTR-DAB2) from PathLinker’s reconstruction of the Wnt signaling pathway and showed that CFTR is associated with Wnt signaling through interactions between RYK, a non-canonical receptor, and DAB2, an inhibitor of canonical Wnt signaling [6].

Despite PathLinker’s success, calculating the first 20,000 shortest paths from receptors to transcriptional regulators for each signaling pathway in the NetPath databases captured only about 70% of the known signaling interactions [6], leaving much room for improvement. Extensions of PathLinker have used auxiliary data such as protein localization information [10] or additional downstream processing [11] to accurately reconstruct ground truth pathways. Other extensions have constrained the paths from *R* to *T* to follow regular expression patterns [12].

### 1.2 Contributions

Inspired by the success of KSP approaches, we reframe the Pathway Reconstruction Problem to directly optimize an objective function related to path cost on directed acyclic graphs (DAGs). We acknowledge up front that cycles (in the form of feedback loops) are important in signaling pathways, so this problem formulation is narrower in scope than the original Pathway Reconstruction Problem. Next, we present two pathway reconstruction methods that solve this new variant of the problem and apply them to six signaling pathways from the NetPath database [3]. We show that DAG reconstructions exhibit different topologies, they often outperform a previous reconstruction method in recovering annotated proteins and interactions, and the predicted nodes are enriched for different biological processes. Finally, we highlight our method’s ability to reconstruct pathways starting from a larger seed DAG, such as a DAG constructed from known pathway interactions.

## 2 Methods

Let the interactome *G* = (*V, E*) be directed with edge weights *w*_*uv*_ for every edge (*u, v*) ∈ *E*. Given *R, T* ⊂ *V* denoting the set of receptors and set of transcription factors for a particular pathway of interest, we first modify the graph by (a) introducing a super source node *s* to *G*, adding directed edges with 0 edge weight from *s* to each node in *R*, and removing all other incoming edges to nodes in *R* and (b) introducing a super sink node *t* to *G*, adding directed edges with 0 edge weight from each node in *T* to *t*, and removing all other outgoing edges from nodes in *T*. We now focus on finding *s*-*t* paths in *G*, which corresponds to finding paths from *any* receptor in *R* to *any* transcriptional regulator in *T* that have the same cost as the original path lengths from receptors to transcriptional regulators. Further, nodes in *R* and *T* will only start and end the paths (they will never be internal nodes on paths). KSP approaches are parameterized by *k*, the number of shortest paths from *s* to *t*. KSP approaches iteratively “grow” a subgraph *G*_*k*_ of *G* by taking the union of the first *k* shortest paths from *s* to *t*: *G*_1_, *G*_2_, …, *G*_*j*_, …, *G*_*k*_. In this way, KSP approaches iteratively “grow” a subgraph of *G*. Our goal is to grow directed acyclic graphs (DAGs) from a graph *G*.

### 2.1 A Problem Formulation for Growing DAGs

We now formulate an optimization problems to grow directed acyclic graphs (DAGs) from a graph *G*. Let *G*_0_ ⊂ *G* be any DAG that connects *s* and *t* (for example, the shortest *s*-*t* path). We will also keep track of all paths in a DAG: let *P*_*j*_(*s, t*) be the set of paths from *s* to *t* in DAG *G*_*j*_. Finally, we define the new edges added to *G*_*j*_ to be *E*_Δ_ = *E*_*j*_ *\ E*_*j*−1_ and the set of new paths added to be *P*_Δ_ = *P*_*j*_(*s, t*) \ *P*_*j*−1_(*s, t*).

#### The Growing DAG Problem

Given a weighted, directed graph *G* = (*V, E, w*) modified with super-source and super-sink nodes *s* and *t*, a DAG *G*_0_ ⊂ *G* that connects *s* and *t*, and a parameter *k*. For *j* = 1, 2, …, *k*, find a DAG *G*_*j*_ = (*V*_*j*_, *E*_*j*_, *w*_*j*_) where *G*_*j*−1_ ⊂ *G*_*j*_ ⊆ *G* such that

1. *P*_Δ_ ≠∅ (there exists at least one new *s*-*t* path).
2. 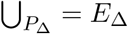 (all new edges in *G*_*j*_ are on some *s*-*t* path).
3. *G*_*j*_ minimizes some cost function *c* : *G*_*j*_ ↦ ℝ, which may be one of the following:

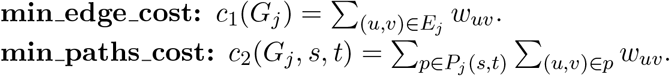

Cost function **min_edge_cost** is simply the cost of the edges in the subgraph *G*_*j*_, while cost function **min_paths_cost** computes the cost of all paths in *G*_*j*_. An example iteration is shown in Figure 1; note that the *G*_1_ that minimizes the two costs functions are quite different in (C) and (D), though they both add at least one new path and all new edges are on some *s*-*t* path.

**Figure 1:**
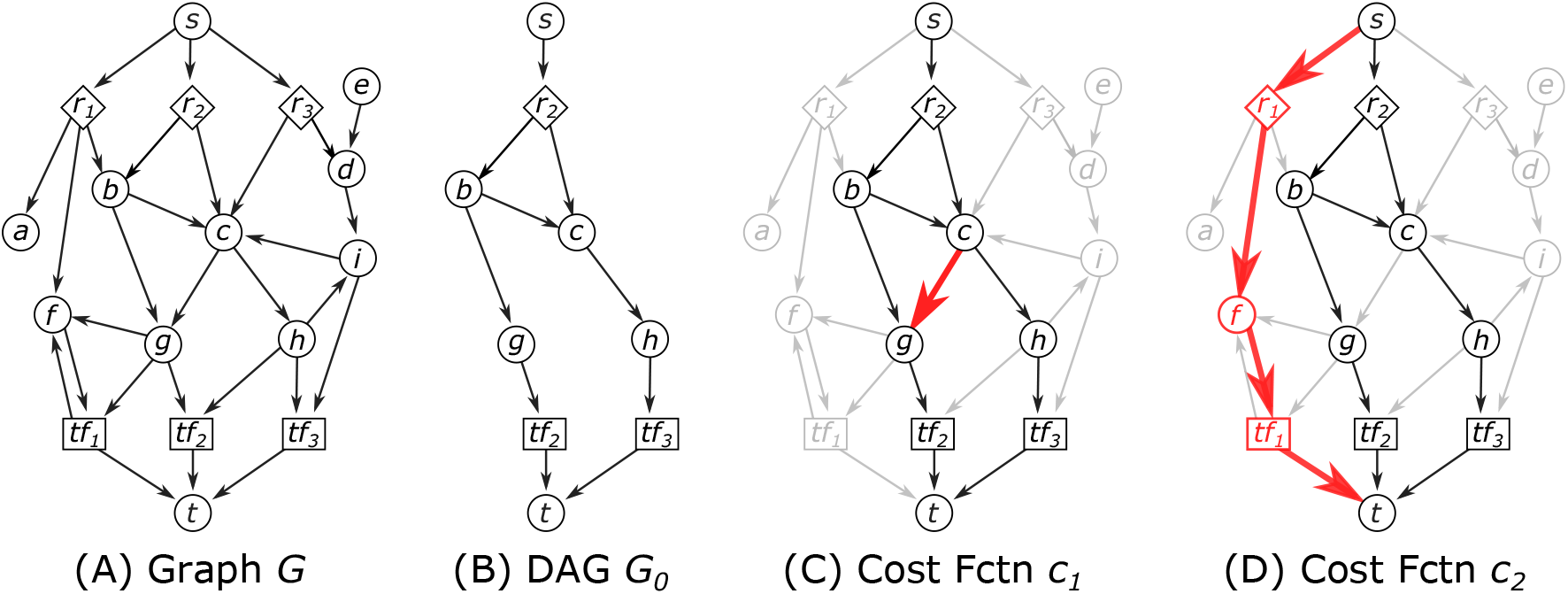
The Growing DAG Problem. (A) Input graph *G* which may contain cycles and bidirected edges. Node *s* is connected to three receptors *r*_1_, *r*_2_, *r*_3_; three transcription factors *tf*_1_, *tf*_2_, *tf*_3_ are connected to node *t*. In this example, all edges have unit cost. (B) An example DAG *G*_0_ ⊂ *G*. (C) An example *G*_1_ that minimizes **min_edge_cost** by adding a single edge to *G*_0_. (D) An example *G*_1_ that minimizes **min_paths_cost** by adding a path of length 4 to *G*_0_.

The shortest-path aspect of this problem comes from minimizing the cost functions. Note that, while KSP approaches will always calculate the *j*th shortest *s*-*t* path in *G*, this path may be comprised of edges already in *G*_*j*−1_ and thus fail the first condition above. KSP approaches are also not guaranteed to produce a DAG at each iteration.

### 2.2 Calculating Cost Functions

In order to implement the Growing DAG Problem, we must show that we can efficiently determine if a DAG *G*_*j*_ satisfies the properties listed and that we can efficiently identify such a DAG from the previous iteration. We first show that, given a DAG *G*_*j*_, we can efficiently calculate the cost functions. Given *G*_*j*−1_ and *G*_*j*_, we can efficiently calculate the cost functions described above. Calculating **min_edge_cost** is straightforward: simply sum the edge weights in *G*_*j*_. However, calculating **min_paths_cost** requires an enumeration of all paths in *G*_*j*_. To calculate **min_paths_cost**, we want to compute the cost of all *s*-*t* paths *P*_*j*_(*s, t*) in *G*_*j*_ without having to enumerate all paths.

It can be rewritten to calculate, for each edge in *E*_*j*_, the number of *s*-*t* paths that contain that edge. Let *f*_*uv*_ be the number of paths in *P*_*j*_(*s, t*) that contain edge (*u, v*):

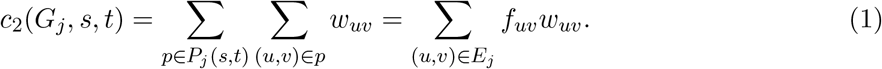

While counting the number of *s*-*t* paths in a general graph is *#P* -complete^1^ [13], we can efficiently count the number of *s*-*t* paths in a DAG. Further, the dynamic program to count the number of *s*-*t* paths in a DAG will also compute the number of *s*-*t* paths *f*_*uv*_ that pass through every edge (*u, v*) ∈ *E*_*j*_. Supplementary Section S1.1 describes the `PathCounter`() algorithm, which returns *f*_*uv*_ for every edge (*u, v*) in a graph. Once we have *f* from the `PathCounter`() algorithm, calculating **min_paths_cost** is straightforward using Equation (1).

### 2.3 Properties of an Optimal *G*_*j*_

Now that we have shown that we can efficiently calculate the cost functions given a DAG *G*_*j*_, we will describe how to identify the possible extensions of *G*_*j*−1_ that guarantee that at least one new *s*-*t* path is added to *G*_*j*_. There may be multiple edges that are added to grow *G*_*j*_ from *G*_*j*−1_; let *G*_Δ_ = (*V*_*j*_ *\ V*_*j*−1_, *E*_*j*_ *\ E*_*j*−1_) be the difference of *G*_*j*_ and *G*_*j*−1_. Since *G*_*j*−1_ and *G*_*j*_ are DAGs, the graph *G*_Δ_ will also be a DAG (which may be disconnected). We first prove some properties of *G*_Δ_ that hold for either of the cost functions listed.

#### Lemma 1.

*Given G*_*j*−1_ *and G*_*j*_ *that satisfy the Growing DAG problem, let G*_Δ_ = *G*_*j*_ *\ G*_*j*−1_. *The G*_Δ_ *that minimizes any of the cost functions will be exactly one path*.

*Proof. G*_Δ_ is a non-empty DAG, since at least one new *s*-*t* path must exist in *G*_*j*_. If *G*_Δ_ is not exactly one path, then *G*_Δ_ must be composed of multiple paths. Since all new edges in *G*_*j*_ must be on some *s*-*t* path, then every maximal path in *G*_Δ_ must start and end on some node from *G*_*j*−1_ (which may include *s* and *t*). Let *P* = *{p*_1_, *p*_2_, …*}* be the set of distinct maximal paths from *G*_Δ_.

If one path *p*_*i*_ ∈ *P* establishes a new *s*-*t* path in *G*_*j*_, then other paths are unnecessary. Let *p*_*i*_ start at some node *v*_0_ and end at some node *v*_*k*_; *p*_*i*_ establishes a new *s*-*t* path if (*s ⇝ v*_0_ ⇝ *v*_*k*_ ⇝ *t*) is in *G*_*j*_. The other paths in *P* will add extraneous edges that are not necessary to establish a new path, which increases cost function **min_edge_cost**. Further, the other paths in *P* will add additional *s*-*t* paths in *G*_*j*_, which increases the cost function **min_paths_cost**. Since maximal paths can be dropped from *P* to minimize either cost function, *G*_Δ_ is not optimal.

Suppose instead that multiple paths are needed to establish a new *s*-*t* path in *G*_*j*_. Without loss of generality, let two paths from *P* be *v*_0_ ⇝ *v*_*k*_ and *u*_0_ ⇝ *u*_*l*_, and let a new *s*-*t* path in *G*_*j*_ be

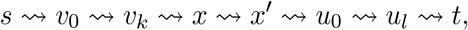

where *x ⇝ x*′ is a path in *G*_*j*−1_. There must be at least one edge from *G*_*j*−1_ between *v*_*k*_ and *u*_0_ because otherwise *p*_*i*_ and *p*_*j*_ would form a single maximal path in *G*_Δ_. Since *G*_*j*−1_ is a connected DAG, then *x ⇝ x′* must be upstream of *t* and/or downstream of *s* in a topological ordering of *V*_*j*−1_. Therefore, at least one of the following paths must exist:

1. *s ⇝ v*_0_ ⇝ *v*_*k*_ *⇝ x ⇝ x*′ ⇝ *t* if *x* is upstream of *t*. In this case, *p*_*i*_ is used but *p*_*j*_ is extraneous and only increases the cost functions (by adding extra edges for **min_edge_cost** and adding extra *s*-*t* paths for **min_paths_cost**), so *G*_Δ_ is not optimal.
2. *s ⇝ x ⇝ x′ ⇝ u*_0_ ⇝ *v*_*l*_ ⇝ *t* if *x* is downstream of *s*. In this case, *p*_*j*_ is used but *p*_*i*_ is extraneous and only increases the cost functions (by adding extra edges for **min_edge_cost** and adding extra *s*-*t* paths for **min_paths_cost**), so *G*_Δ_ is not optimal.□

#### Lemma 2.

*G*_Δ_ *that minimizes the cost functions defines a path that starts at some node in G*_*j*−1_ *and ends at some node in G*_*j*−1_. *All internal nodes on the path are in G*_*j*_ *but not in G*_*j*−1_.

*Proof. G*_Δ_ is a single path *p* = (*v*_0_, *v*_1_, …, *v*_*k*_) by Lemma 1. We need to show that *v*_0_ and *v*_*k*_ are in *V*_*j*−1_, and all other nodes are not in *V*_*j*−1_. First observe that there must be at least two nodes in *G*_Δ_ that are in *G*_*j*−1_ for an *s*-*t* path to include new edges in *G*_*j*_ (which may include *s* and/or *t*); call these nodes *x* and *y*. Consider the path in *G*_*j*_ *s ⇝ x ⇝ y ⇝ t* where *x ⇝ y* is a path in *G*_Δ_.

1. There are exactly two nodes *x, y* ∈ *V*_*j*−1_ in the path *p* = (*v*_0_, *v*_1_, …, *v*_*k*_). Suppose there was a third node, *z* ∈ *V*_*j*−1_, in the path such that *x ⇝ z ⇝ y*. Since *G*_*j*−1_ is a connected DAG, then *z* must be upstream of *t* and/or downstream of *s*. Therefore, at least one of the following paths must exist:
  a. *s ⇝ x ⇝ z ⇝ t* if *z* is upstream of *t*. In this case, *z ⇝ y* is extraneous and increases the cost functions (by adding extra edges for **min_edge_cost** and adding extra *s*-*t* paths for **min_paths_cost**), so *G*_Δ_ is not optimal.
  b. *s ⇝ z ⇝ y ⇝ t* if *z* is downstream of *s*. In this case, *x ⇝ z* is extraneous and increases the cost functions (by adding extra edges for **min_edge_cost** and adding extra *s*-*t* paths for **min_paths_cost**), so *G*_Δ_ is not optimal.
2. *x* = *v*_0_. Suppose *x* is some node on the path *p* that is not *v*_0_; call it *v*_*i*_. The path from (*v*_0_, …, *v*_*i*−1_) could be dropped, because *s ⇝ x* will bypass these nodes.
3. *y* = *v*_*k*_. Same argument as for *x* = *v*_0_.

□

Together, Lemmas 1 and 2 prove the following theorem:

#### Theorem 1.

*Given G*_*j*−1_ *and G*_*j*_ *that satisfy the Growing DAG problem, G*_*j*−1_ *and G*_*j*_ *differ by exactly one path that starts and ends in G*_*j*−1_ *and contains no other nodes from G*_*j*−1_.

### 2.4 The Growing DAG Algorithm

At each iteration *j*, we keep track of candidate paths for DAG *G*_*j*_ using a modified Dijkstra’s algorithm. We rely on topologically sorting the nodes in a DAG, which results in a partial ordering of the nodes. Let *σ*(*G*) denote the partial ordering of a DAG *G*, with the position of each node *v* denoted by *σ*_*v*_. For a node *v* in a DAG, its *ancestors* are all nodes *u* where *σ*_*u*_ *< σ*_*v*_ and its *descendants* are all nodes *x* where *σ*_*v*_ *< σ*_*x*_). Note that there may also be nodes *y* that is neither an ancestor nor a descendant of *v* (e.g., where *σ*_*v*_ = *σ*_*y*_); we say that *v* and *y* are *incomparable*.

Algorithm 1 takes as input a directed, weighted graph *G*, an initial DAG *G*_0_, the number of iterations *k*, and a cost function *c* (either **min_edge_cost** or **min_paths_cost**). It returns a list of length *k* that denotes the min-cost paths for each iterations according to the specified cost function. To track the set of candidate paths, we use a `dist` dictionary that stores the cost of the best paths between topologically sorted nodes in the DAG. We assume that we also have the predecessors so we can determine the path *u ⇝ v* from `dist`[*u*][*v*] as well.

Lines 3–8 initialize the `dist` dictionary for the nodes in *G*_0_. First, we build a *G*_cand_ graph that removes the existing DAG from *G*. Then, for every node *u* in *G*_0_ we consider, we must also remove any node that is an ancestor of *u* to prevent paths that would induce cycles. Once we have the graph, we call a slightly modified Dijkstra’s algorithm called `MultiTargetDijkstra`() to find the shortest path from a source node *s* to each target node, ensuring that a node in *G*_*j*_ is never considered an internal node on a path and returning early if all target nodes are reached (Supplementary Section S1.2).

Once the distances are initialized, we iteratively grow *G*_*j*_ for *j* from 1 to *k*. First, we identify the min-cost path from `dist` according to cost function *c* (Lines 10–21). This is simple for **min_edge_cost**, when we select the path *u ⇝ v* with the lowest cost in `dist`. For **min_paths_cost**, we need to calculate the cost of all *s*-*t* paths for every possible choice of valid *u ⇝ v* path in `dist`. We can call the `PathCounter`() algorithm (Supplementary Algorithm S1) on a graph *G*^*t*^ that contains the path we are checking in order to calculate the all-paths-cost *C*[*u*][*v*] (Lines 16–18). We select the path *u ⇝ v* with the lowest all-paths-cost *C*[*u*][*v*], add the selected path to *G*_*j*−1_ to make the *G*_*j*_ DAG, and update the distances dictionary.

We implemented a number of speedups to improve the runtime of Algorithm 1, two of which are notable. First, in the `MultiTargetDijkstra`() function call, we can set the target nodes *T* to be only the nodes *v* for which *u ⇝ v* needs to be recalculated (Supplementary Section S1.2). In practice, this dramatically reduced the number of target nodes that `MultiTargetDijkstra`() needed to find, re-running between 0.2%–14% of the possible number of target nodes across all runs. A second speedup comes from avoiding re-computations when minimizing **min_paths_cost**. Lines 16–18 generate a new DAG *G′*, calls `PathCounter`() on this new graph, and uses these values to compute the **min_paths_cost** for every ordered node pair *u, v*. In practice, we first calculate *f* for *G*_*j*_ as the first step in the main `for` loop, and use functions to update *f* as if we have added the path *u ⇝ v* (without needing to create a new graph *G′*). These update functions, similar to determining the `upstream` and `downstream` dictionaries in the `PathCounter`() algorithm, adjusts all paths from *s* to *u* and all paths from *v* to *t* assuming that path *u ⇝ v* has been added.

## 3 Results

We ran `GrowDAGs` for both cost functions to reconstruct six diverse signaling pathways from the NetPath database [3]. We identified receptors and transcriptional regulators for each pathway using previously established resources of these proteins types [14, 15, 16]. The six networks range in size, density, and the number of receptors and transcriptional regulators (Table 1). We used an interactome *G* with 17, 513 nodes and 577, 617 directed edges weighted by experimental evidence [10]. This interactome is built from a combination of molecular interaction data (BioGrid, DIP, InnateDB, IntAct, MINT, PhosphositePlus) and annotated signaling pathway databases (KEGG, NetPath, and SPIKE). Since NetPath nodes and edges are part of the interactome, we can use these pathways as ground truth networks to assess reconstruction methods (similar to the analyses in previous work [6, 10]). The edge weights are negative log transformed to become edge costs, which we aim to minimize.

**Table 1:** Signaling pathways chosen for this study [3]. TRs: transcriptional regulators.

| Name | Abbrev. | Nodes | Edges | Receptors | TRs |
| --- | --- | --- | --- | --- | --- |
| B Cell Receptor Pathway | BCR | 137 | 456 | 1 | 18 |
| Epidermal Growth Factor Receptor Pathway | EGFR1 | 231 | 1456 | 6 | 33 |
| Interleukin 1 Pathway | IL1 | 43 | 178 | 3 | 5 |
| T Cell Receptor Pathway | TCR | 154 | 504 | 7 | 20 |
| Transforming Growth Factor $\beta$ Pathway | TGFβ | 209 | 863 | 5 | 77 |
| Wnt Pathway | Wnt | 106 | 428 | 14 | 14 |

### Algorithm 1 GrowDAGs(*G* = (*V,E*), *G*_0_ = (*V*_0_,*E*_0_), num iters *k*, cost function *c*)

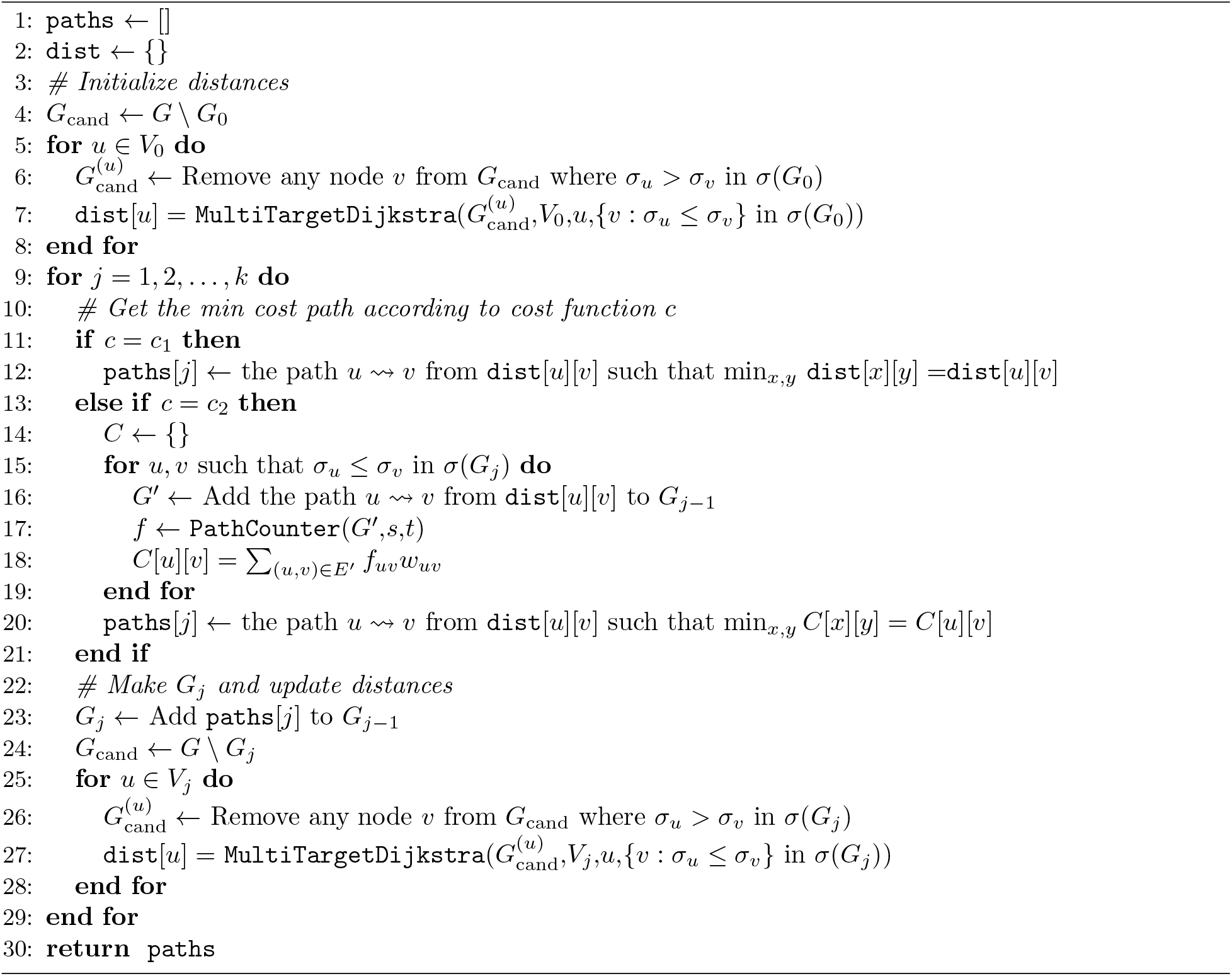

### 3.1 Topological Differences between DAG Reconstructions

We first provide some examples of reconstructed pathways using `GrowDAGs` by examining the topological differences between DAG reconstructions of the Wnt signaling pathway. We grew DAGs from each pathway-specific *G*_0_ defined as the shortest *s*-*t* path in *G*, where super source node *s* is connected to the pathway’s receptors and the pathway’s transcriptional regulators are connected to super sink *t* (Figure 1A). As expected, **min_edge_cost** tended to produce DAGs whose paths reuse the same nodes while **min_paths_cost** tended to produce DAGs with more non-overlapping *s*-*t* paths (Figure 2 shows *k* = 75 for the Wnt pathway reconstructions). Network visualizations of all pathways up to *k* = 200 are available on GraphSpace [17].^2^

**Figure 2:**
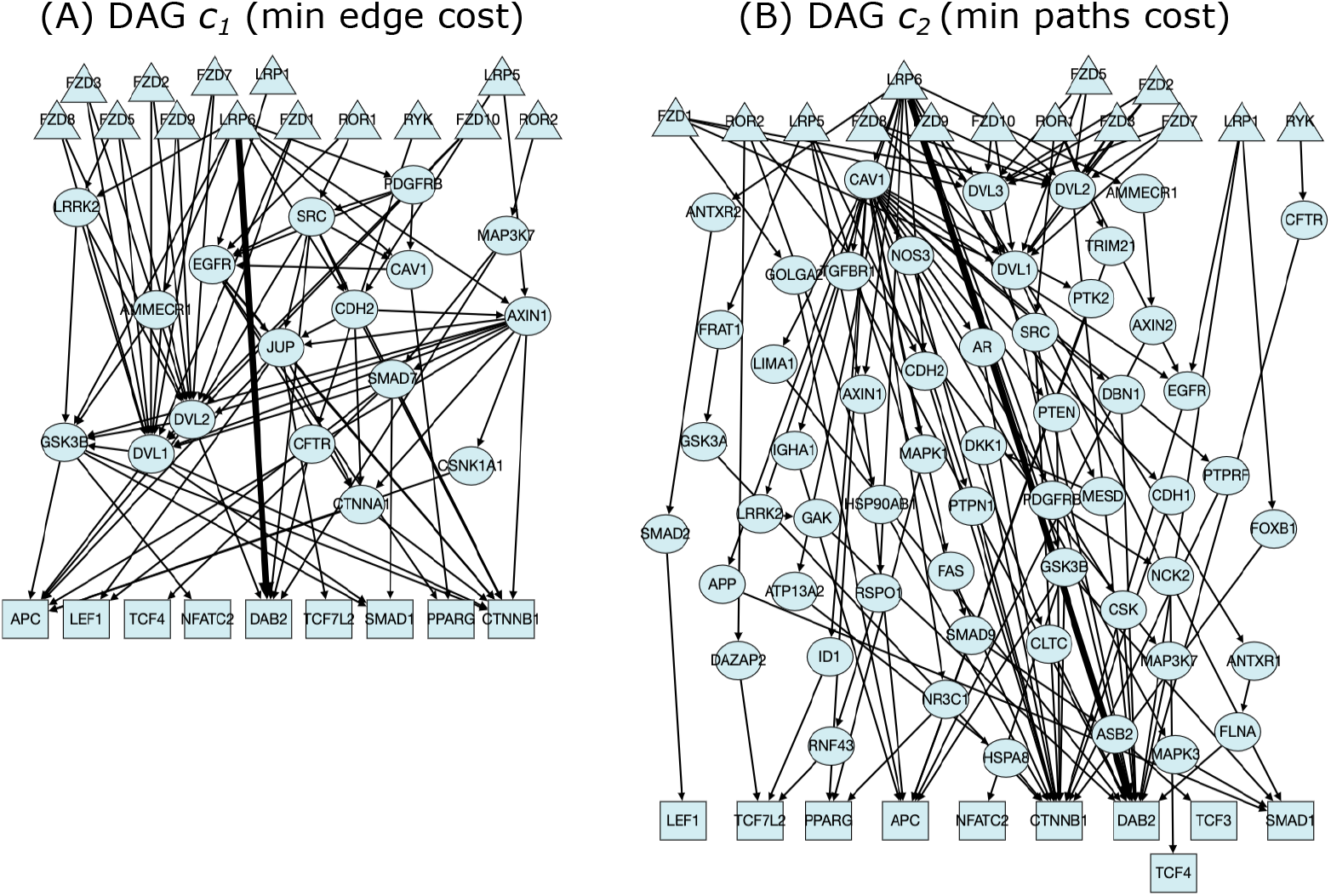
Wnt pathway `GrowDAGs` reconstructions (*k* = 75) for (A) **min_edge_cost** and (B) **min_paths_cost**. The shortest path from any receptor to any transcriptional regulator was used as *G*_0_ (shown as the thick edge).

We also compared the resulting `GrowDAGs` reconstructions with those from PathLinker, the *k*-shortest paths approach [6]. We ran the `GrowDAGs` methods (starting from the shortest-path *G*_0_) and PathLinker on each of the six pathways until the reconstruction contained the same number of nodes or edges as the ground truth pathway (or until *k* = 1000). We call these *size-matched reconstructions*, and they depend on whether the reconstructions are matched by the number of nodes or the number of edges. Supplementary Table S1 shows the *k* values needed for size-matched reconstructions for all six pathways; two node-based reconstructions and three edge-based reconstructions from the BCR pathway and the EGFR1 pathway did not reach the ground truth number of nodes or edges by *k* = 1000, so these comparisons are not necessarily size-matched.

The Wnt size-matched reconstruction for **min_paths_cost** took fewer iterations than **min_edge_cost** or PathLinker for both nodes and edges, indicating that **min_paths_cost** tends to add new nodes and edges to reconstructions rather than reusing existing nodes and edges (Figure 3A). This observation about **min_paths_cost** holds across size-matched reconstructions for all six pathways, though Wnt tends to be an outlier when comparing **min_edge_cost** to PathLinker (Supplementary Figures S2–S3). For all other pathway reconstructions, **min_edge_cost** takes the most iterations to reach size-matched reconstructions for nodes, and PathLinker falls between the two DAG methods.

**Figure 3:**
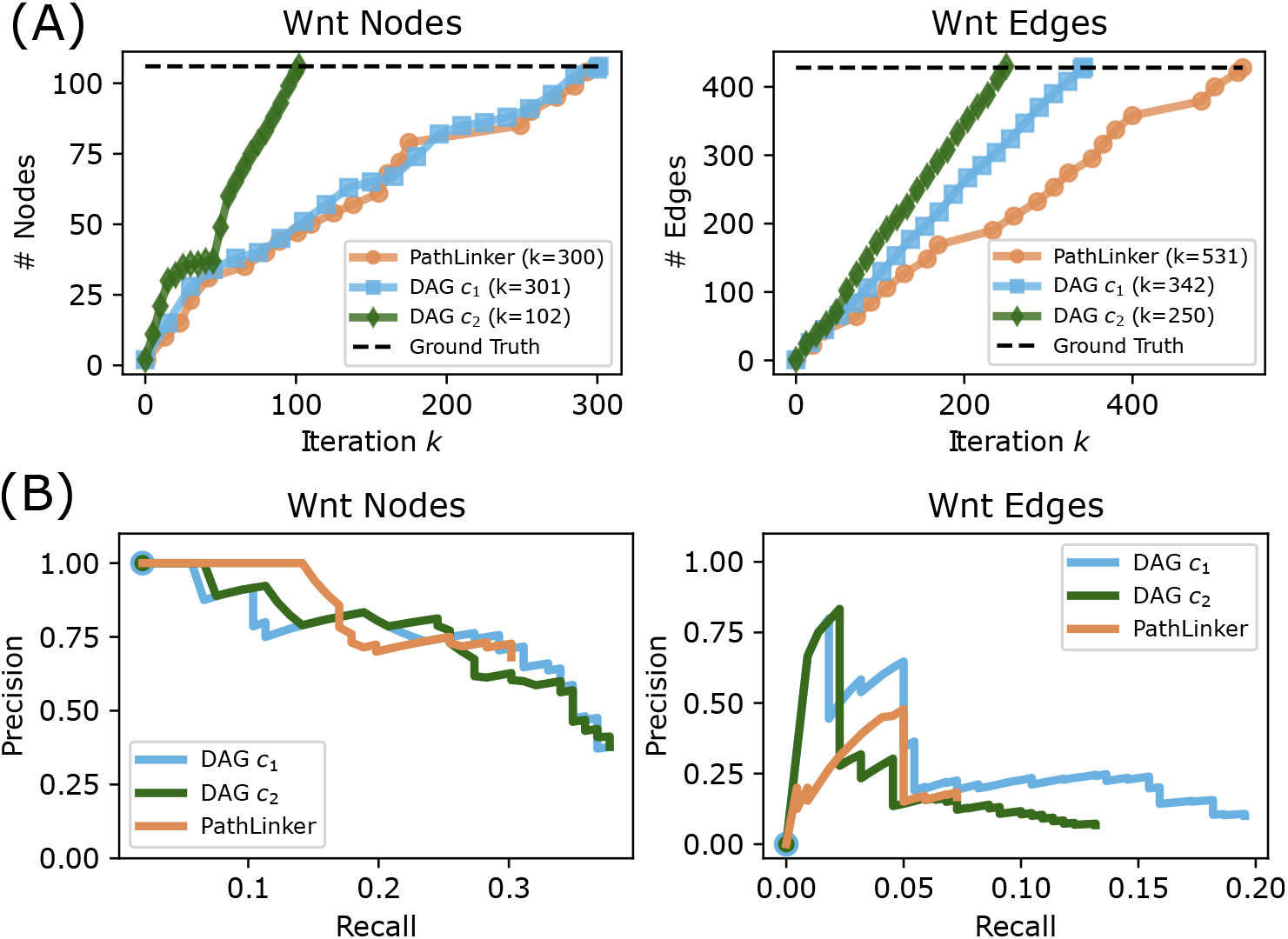
(A) Number of nodes and edges at each iteration of the Wnt pathway reconstructions. Horizontal dashed line indicates the number of nodes and edges in the Wnt NetPath pathway. (B) Precision-recall curves for size-matched reconstructions from the Wnt pathway. *c*_1_: **min_edge_cost**; *c*_2_: **min_paths_cost**.

### 3.2 Comparison to Ground Truth Pathways

We then evaluated how well the `GrowDAG` pathways captured annotated proteins and interactions from the six NetPath pathways. For this analysis, we considered the NetPath pathways ground truth nodes and edges, though it is widely acknowledged that signaling pathway databases are incomplete. For each of the six pathways, we calculated the precision and recall for size-matched reconstructions for nodes and edges (Figure 3B and Supplementary Figures S4–S5). When computing precision and recall of edges, we compare undirected edges to undirected ground truth edges, which is consistent with previous evaluations [6]. Overall, the methods achieve relatively low recall (e.g., 0.15–0.5 recall for nodes) despite relatively high precision (Supplementary Figures S4–S5). The `GrowDAG` reconstructions have a higher AUPRC than PathLinker in all but one case (TCR), whose max AUPRC is an order of magnitude smaller than any other case (Table 2). Of the `GrowDAG` cost functions, **min_paths_cost** has the highest AUPRC for seven cases compared to four cases for **min_edge_cost**.

**Table 2:** Area under the precision-recall (AUPRC) values for nodes (left) and edges (right) for size-matched reconstructions. *c*_1_: **min_edge_cost**; *c*_2_: **min_paths_cost**. Largest AUPRC values are shown in bold; asterisks denote reconstructions that did not reach the number of ground truth nodes or edges by *k* = 1000 (see Supplementary Table S1).

| Name | Node AUPRC |  |  | Edge AUPRC |  |  |
| --- | --- | --- | --- | --- | --- | --- |
| | DAG $c_1$ | DAG $c_2$ | PL | DAG $c_1$ | DAG $c_2$ | PL |
| BCR | 0.179 | **0.242\*** | 0.132 | 0.018 | **0.029** | 0.003 |
| EGFR1 | **0.275** | 0.249* | 0.234 | **0.132\*** | 0.013* | 0.012* |
| IL1 | 0.148 | **0.259** | 0.225 | 0.020 | **0.047** | 0.030 |
| TCR | 0.130 | **0.187** | 0.176 | 0.003 | 0.003 | **0.006** |
| TGFβ | **0.397** | 0.377 | 0.355 | 0.052 | **0.060** | 0.036 |
| Wnt | 0.274 | **0.276** | 0.272 | **0.057** | 0.030 | 0.028 |

While the methods have low recall, overall size matched-sized reconstructions are often too large for visual exploration (for example, the networks visualized in Figure 2 are only a subset of the matched-sized reconstructions). Instead, considering the precision for the top 50 predictions (e.g., the first 50 nodes or first 50 edges in a reconstruction) is a complementary assessment in terms of reconstruction usefulness. When considering the first 50 predictions, DAG **min_paths_cost** no longer consistently outperforms the other two methods, though one of the `GrowDAG` reconstructions has the highest precision for five of the node reconstructions and five of the edge reconstructions (Table 3).

**Table 3:**
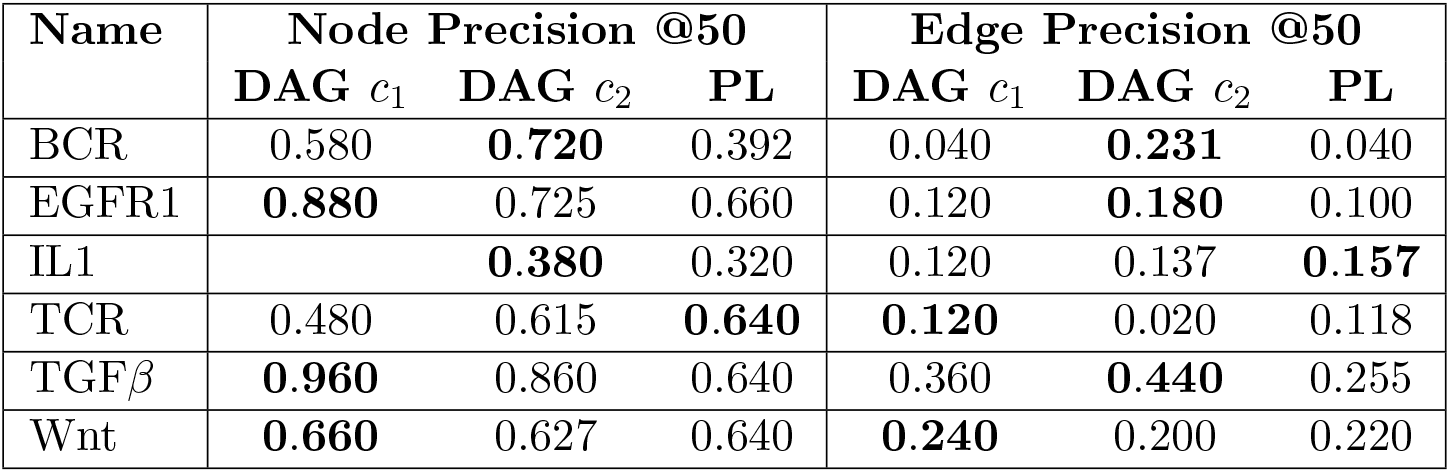
Precision of the first 50 predictions (nodes on the left, edges on the right) for the reconstructions. *c*_1_: **min_edge_cost**; *c*_2_: **min_paths_cost**. Largest values are shown in bold; missing entries denote that 50 unique predictions weren’t reached at the specified value of *k* for sized-matched reconstructions.

| Name | Node Precision @50 |  |  | Edge Precision @50 |  |  |
| --- | --- | --- | --- | --- | --- | --- |
| | DAG $c_1$ | DAG $c_2$ | PL | DAG $c_1$ | DAG $c_2$ | PL |
| BCR | 0.580 | **0.720** | 0.392 | 0.040 | **0.231** | 0.040 |
| EGFR1 | **0.880** | 0.725 | 0.660 | 0.120 | **0.180** | 0.100 |
| IL1 |  | **0.380** | 0.320 | 0.120 | 0.137 | **0.157** |
| TCR | 0.480 | 0.615 | **0.640** | **0.120** | 0.020 | 0.118 |
| TGFβ | **0.960** | 0.860 | 0.640 | 0.360 | **0.440** | 0.255 |
| Wnt | **0.660** | 0.627 | 0.640 | **0.240** | 0.200 | 0.220 |

One of the reasons for the low precision, especially for edges, is that the number of negative examples is vastly larger than the number of positive examples for a single pathway. Thus, previous evaluation frameworks have subsampled the negative examples from the interactome to ensure a 50:1 ratio of negatives to positives [6]). This inflates the precision but can also more clearly separate performances of different approaches. We conducted the same precision-recall analysis by subsampling negatives in a 50:1 ratio and repeating for 10 iterations – the `GrowDAG` reconstructions outperform PathLinker even when subsampling negatives, though more **min_edge_cost** reconstructions have the highest AUPRC across methods (Supplementary Figures S6–S7 and Table S2).

### 3.3 Reconstructions Capture Diverse Biological Processes

Next, we evaluated whether the three pathway reconstruction methods recover different biological processes. Here, we considered the nodes in the sized-matched reconstructions and found that, for five of the six pathways, each method contains distinct nodes not found in the other reconstructions (Supplementary Figure S8). For example, only 17% of the predicted nodes for Wnt size-matched reconstructions are predicted by all methods (Figure 4). We performed gene function enrichment using PantherDB [18, 19] on the uniquely-predicted nodes for each method (e.g., 54 DAG **min_edge_cost** nodes, 46 DAG **min_paths_cost** nodes, and 48 PathLinker nodes in Figure 4) to determine whether any Panther pathways are enriched in each of these unique sets. Specifically, we ran the PANTHER Over-representation Test (version 17.0) using Fisher’s exact test and an FDR correction of 0.05.

**Figure 4:**
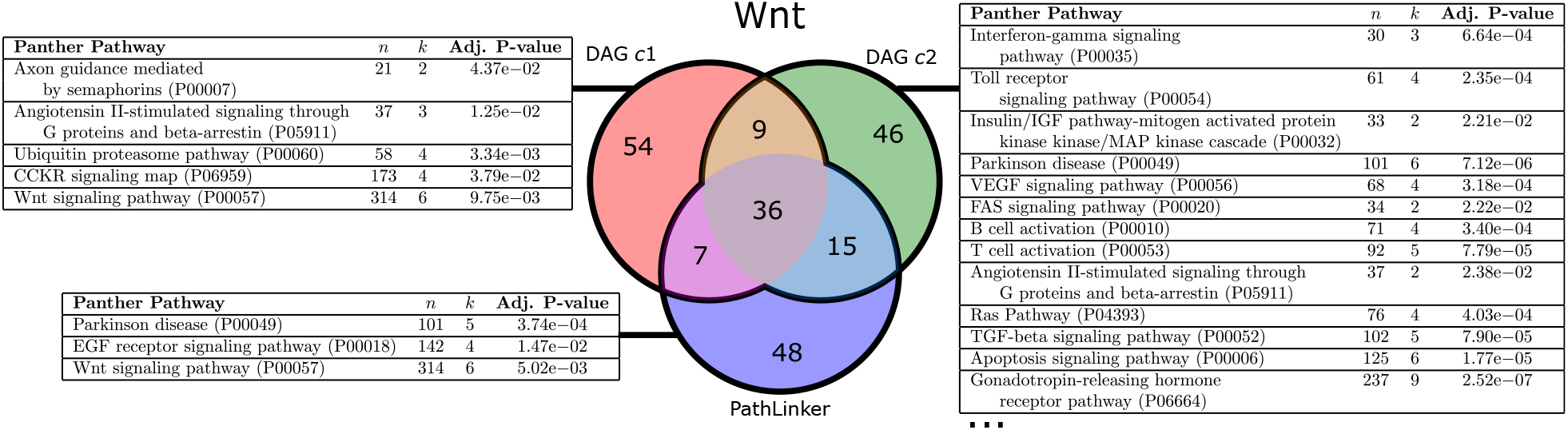
Venn diagram of predicted nodes for `GrowDAG` methods and PathLinker for size-matched Wnt reconstructions. Tables show gene function enrichment of uniquely-predicted nodes using PantherDB pathways. *n*: number of proteins in the Panther pathway; *k*: number of predicted proteins in the Panther pathway; c1: **min_edge_cost**; c2: **min_paths_cost**. Full table for DAG **min_paths_cost** is shown in Supplementary Table S3.

We found that each set of unique nodes were enriched for at least three Panther pathways (Tables in Figure 4). DAG **min_paths_cost** has 24 enriched Panther pathways; the full table is in Supplementary Table S3. All three reconstructions contain nodes that are enriched in Wnt signaling (DAG **min_paths_cost** captures three nodes with an adjusted *p*-value of 3.62 *×* 10^−2^, Supplementary Table S3), each capturing different aspects of the Wnt pathway that is separate from the other two methods. DAG **min_paths_cost** and PathLinker reconstructions are also enriched for Parkinson Disease, which has a known connection to Wnt signaling through inflammatory pathways [20]. DAG **min_paths_cost** also appears to be enriched in many signaling pathways that are known to cross-talk with Wnt, including T cell signaling [21], B cell signaling [22], and EGFR signaling [23], among others.

### 3.4 *G*_0_ as NetPath DAGs

One of the powerful aspects of `GrowDAGs` is the ability to begin the reconstruction with any DAG. For example, using the ground truth pathways as input DAGs, our methods will propose new nodes and edges that are not currently annotated to that pathway. To illustrate this potential, we converted each ground truth pathway from NetPath into a *G*_0_ DAG by considering only the nodes and edges in the ground truth network (Table 1) after connecting super source node *s* to all receptors and super sink node *t* to all transcriptional regulators (similar to the modifications made to the full interactome). We calculated up to 1, 000 shortest *s*-*t* paths in the ground truth network (e.g., by running PathLinker) and built *G*_0_ by adding each path only if it did not introduce a cycle in *G*_0_. Since the NetPath edges are used to build the interactome, each ground truth DAG conversion *G*_0_ is, by construction, a subgraph of the interactome. The total number of nodes and edges in the DAG conversion of *G*_0_ were correlated with the overall size of the ground truth pathway (Table 4). We ran `GrowDAGs` for *k* = 25 in order to be able to visualize the newly-added nodes and edges.

**Table 4:** The ground truth pathways converted to DAGs, which are used as *G*_0_ in Section 3.4. The total number of PathLinker paths are also reported.

| Name | PathLinker<br>Paths | Nodes in<br>DAG Conversion | Edges in<br>DAG Conversion |
| --- | --- | --- | --- |
| BCR | 958 | 58 | 131 |
| EGFR1 | >1000 | 105 | 461 |
| IL1 | 713 | 22 | 55 |
| TCR | 952 | 57 | 139 |
| TGFβ | 70 | 33 | 70 |
| Wnt | 984 | 40 | 89 |

Running `GrowDAGs` on the six signaling pathways with the ground truth DAGs *G*_0_ produced similar trends in network topology for **min_edge_cost** and **min_paths_cost** that were shown in Section 3.1. For the 25 new paths added to the NetPath DAGs, we also counted the number of directed edges that were present in Netpath or KEGG pathway databases, indicating that a known signaling interaction was recovered. We found that, on average, 16.9 *±* 3.4 directed edges from DAGs generated by **min_edge_cost** and 13.2 *±* 3.0 directed edges from DAGs generated by **min_paths_cost** were present in the databases. An example reconstruction for IL1, the smallest pathway, is shown in Figure 5. These reconstructions potentially suggest new proteins and interactions that should be considered in the annotated pathways, and the percentage of interactions from signaling pathway databases are a promising sign of identifying signaling edges in these DAGs.

**Figure 5:**
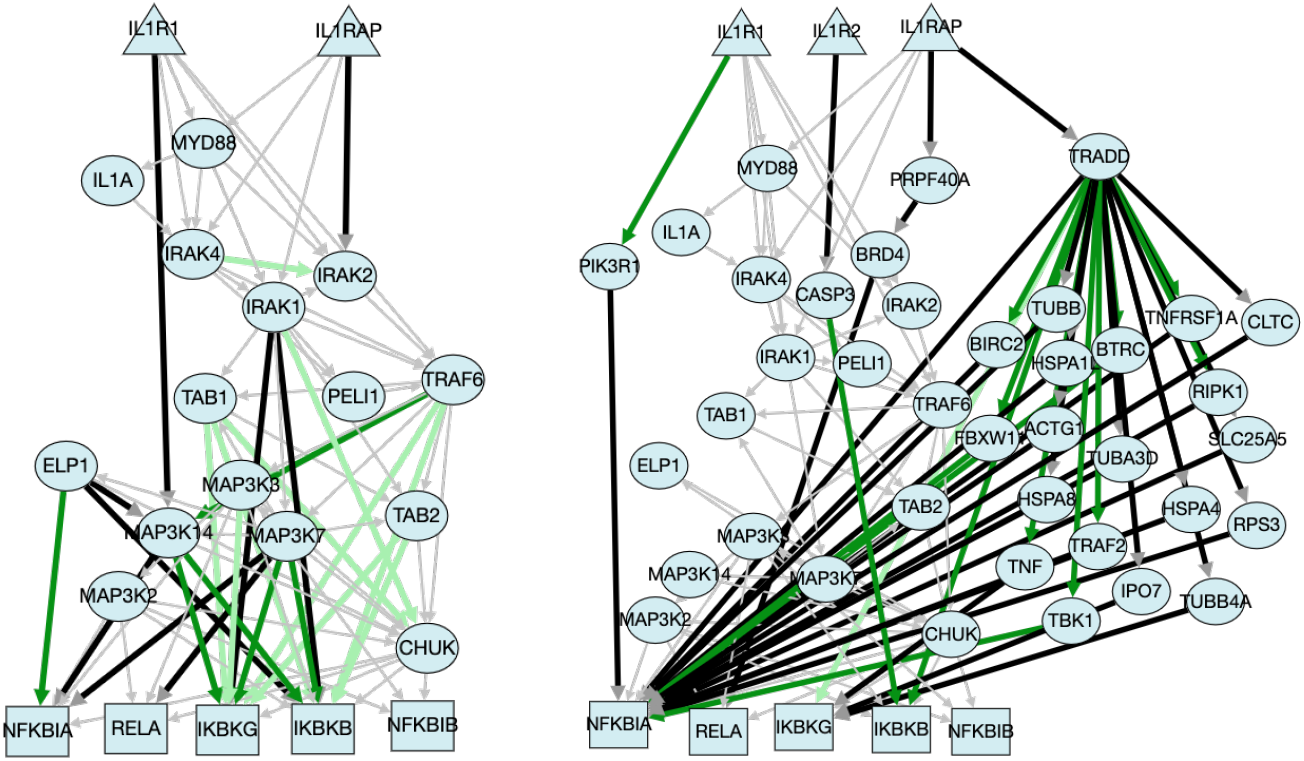
IL1 Reconstructions for (left) **min_edge_cost** and (right) **min_paths_ost** for *k* = 25. Here, *G*_0_ is shown in thin gray edges. Dark green edges appear in the NetPath database; light green edges appear in the KEGG database.

## 4 Discussion

We have described the Growing DAG Problem, a pathway reconstruction formulation that directly adds paths to a DAG that explicitly optimize one of two cost functions (minimizing edge costs or minimizing *s*-*t* path costs). We presented the `GrowDAGs` algorithm that iteratively finds the most optimal DAG *G*_*j*_. Applying this algorithm to six diverse NetPath pathways, we show that **min_edge_cost** and **min_paths_cost** admits DAGs with different network topologies that often outperforms a KSP approach compared to the annotated proteins and interactions in NetPath. Further, the reconstructions return proteins that are enriched for different biological processes. We also demonstrate that we can begin with a larger DAG *G*_0_ such as a DAG converted from the NetPath ground truth. To our knowledge, this is the first pathway reconstruction algorithm that can grow a network based on a seeded subnetwork (rather than a set of nodes).

In our analysis of `GrowDAGs` and PathLinker reconstructions from six pathways, the `GrowDAGs` methods outperformed PathLinker for size-matched reconstructions in all scenarios except one. Between the two cost functions, DAG **min_paths_cost** had a higher AUPRC for more scenarios than DAG **min_edge_cost** (Table 2). It is worth noting, however, that DAG **min_edge_cost** performed the best on the EGFR1 reconstructions, which is the largest ground truth network we tested. DAG **min_edge_cost** also performed better than the other reconstructions for TFG*β* when reconstructing size-matched nodes (Table 2). Both EGFR1 and TGF*β* node reconstructions have high overlap between DAG **min_paths_cost** and PathLinker (as evidenced by the venn diagrams in Supplementary Figure S8). This suggests that, when DAG **min_paths_cost** and PathLinker produce similar reconstructions, DAG **min_edge_cost** may better reflect nodes in the ground truth pathway.

The `GrowDAG` formulations rely on a parameter *k*, rely on a parameter *k*, which determines how large to grow the pathway reconstruction. The choice of *k* dramatically changes the size of the reconstructions, and this user-defined parameter should be chosen carefully with the user’s goal in mind. For example, we calculated up to *k* = 1000 iterations in order to assess how well entire pathways could be reconstructed (though pushing this to larger values of *k* will have challenges, as described below). However, when looking for potentially new proteins and interactions that are missing from an annotated pathway, the first few predictions (e.g., the top 50 predictions shown in Table 3) may be more appropriate. For visualization purposes, *k* = 200 begins to reach the limit of useful network visualization (see, for example, the reconstructions available at http://graphspace.org/graphs/?query=tags:DAG#). In general, larger values of *k* will provide a better sense of how much of the pathway is captured in a reconstruction, whereas looking at the top predictions for small values of *k* will provide a ranked list of potentially new proteins and interactions that are missing from the annotated pathway.

The Growing DAG Problem and subsequent algorithms are limited by the DAG constraints – namely, that reconstructions will never contain cycles. Feedback loops are an integral part of signaling [24], and many of the signaling pathways in NetPath, KEGG, and other pathway databases have documented examples of these cycles. Our reasons for starting with DAGs were twofold: first, intracellular signaling that begins at a cell membrane and ends with transcriptional regulation has a general direction; and second, DAGs provide a reduced search space when growing networks. Beyond the search space for optimal subgraphs at each iteration, the **min_paths_cost** optimization criterion could not be efficiently computed on a subgraph with cycles, though there may very well be another optimization function that would be useful to consider in general graphs. Generalizing the computational problem and the general framework to graphs with cycles would be an important future step in pathway reconstruction.

While the `GrowDAGs` framework is a promising way to reconstruct signaling pathways with certain topologies, a few limitations remain that prohibit its widespread use for large pathway reconstructions. First, the algorithm is still slow in practice, namely due to having to check all pairs of topologically sorted nodes of *G*_*j*−1_ at each iteration. Runtimes for *k* = 100 averaged 2.1 hours, ranging from eleven minutes to 16 hours (a full table of runtimes is available in Supplementary Table S4). However, computing the next iterations for *k* = 101, …, 200 averaged 19.6 hours, ranging from 44 minutes to nearly 52 hours. While we mention a few speedups in Section 2.4, more improvements are needed to reconstruct pathways at the scale of PathLinker (which calculated reconstructions up to *k* = 20, 000). Second, other cost functions may produce different topologies than those reconstructed here; for example, minimizing the average cost of new paths or taking a weighted average of **min_edge_cost** and **min_paths_cost**. However, Lemma 2 does not hold for such cost functions.

Despite these challenges, the Growing DAG Problem presents a first step towards explicitly optimizing a cost function related to pathway reconstruction. Existing pathway reconstruction methods will always return networks with the same topologies – *k* shortest paths, Steiner forests, random walks, etc. – and the proposed cost functions provide control over the topologies of the `GrowDAG` reconstructions. This work opens the door for improved cost functions that reflect ground truth pathways and provides a framework for designing algorithms to grow networks.

## Data and Code Availability

All code and data related to Growing DAGs are available on GitHub: https://github.com/Reed-CompBio/growing-dags. Pathway reconstruction visualizations for the six pathways and *k* = 200 are available on GraphSpace [17]: http://graphspace.org/graphs/?query=tags:DAG#.

## Acknowledgements

This work was supported by the NSF (DBI-1750981) to AR. We thank Layla Oesper, Ibrahim Youssef, and Tobias Rubel for feedback on early versions of this manuscript.

## Footnotes

1 #P (Sharp-P) problems are the counting problems associated with decision problems in *NP*.

2 http://graphspace.org/graphs/?query=tags:DAG#.

