## Supplementary Information for "Growing DAGs: Optimization Functions for Pathway Reconstruction Algorithms"

### S1 Supplementary Methods

#### S1.1 The `PathCounter()` Algorithm

To calculate **min_paths_cost**, we want to compute the cost of all $s$-$t$ paths $P_j(s,t)$ in a DAG $G_j$ without having to enumerate all paths. The `PathCounter`() algorithm is a dynamic program that computes the number of $s$-$t$ paths $f_{uv}$ that pass through every edge $(u,v)$ in the DAG. Algorithm S1 relies on topologically sorting the nodes in the DAG $G$. For every node $v \in G_j$, we can decompose the $s$-$t$ paths that go through $v$ into paths from $v$ to $t$ (Lines 2–5) and from $s$ to $v$ (Lines 6–9). In order to compute $f_{uv}$, we must calculate the number of $s$-$t$ paths that go through some edge $(u,v)$. This value can be calculated by multiplying `upstream[v]` and `downstream[u]` ((Lines 10–12)). An illustrative example is shown in Figure S1.

**Algorithm S1** `PathCounter`(DAG $G = (V,E)$,$s$,$t$)

1: `nodes` $\leftarrow$ topologically sort $V$
2: `upstream` $\leftarrow \{\}$
3: **for** $u$ in `reverse`(`nodes`) **do**
4: $\quad$ `upstream`$[u] \leftarrow \begin{cases} \sum_{(u,v)\in E} \texttt{upstream}[v] & \text{if there exists some outgoing edge from } u \\ 1 & \text{if } u = t \\ 0 & \text{otherwise} \end{cases}$
5: **end for**
6: `downstream` $\leftarrow \{\}$
7: **for** $v$ in `nodes` **do**
8: $\quad$ `downstream`$[v] \leftarrow \begin{cases} \sum_{(u,v)\in E} \texttt{downstream}[u] & \text{if there exists some incoming edge to } v \\ 1 & \text{if } v = s \\ 0 & \text{otherwise} \end{cases}$
9: **end for**
10: **for** $(u,v) \in E$ **do**
11: $\quad f_{uv} = \texttt{upstream}[v] \times \texttt{downstream}[u]$
12: **end for**
13: **return** $f$

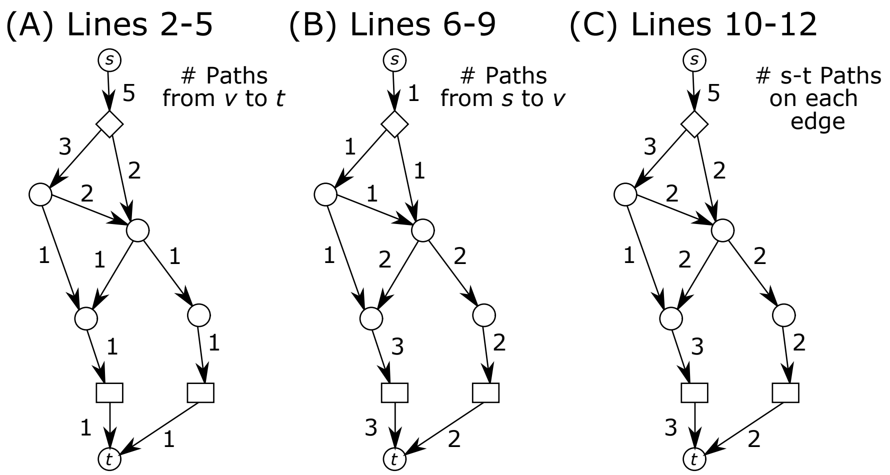

Figure S1: Illustration counting the total number of $s$-$t$ paths by (A) computing the number of paths from $v$ to $t$, (B) computing the number of paths from $s$ to $v$, and (C) combining these values to get the number of times each edge is traversed to count the total number of $s$-$t$ paths.

#### S1.2 The `MultiTargetDijkstra()` Algorithm

After building a candidate graph for a node $u$ (e.g., $G_{\text{cand}}^{(u)}$ in Algorithm 1 of the main text), we wish to calculate the distances from $u$ to $u$'s descendants and incomparable nodes from $\sigma(G_j)$:

$$T = \{v : \sigma_u \leq \sigma_v\} \text{ in } \sigma(G_j).$$

We call a slightly modified Dijkstra's algorithm called `MultiTargetDijkstra()` to find the shortest path from $u$ to each target node in $T$ (Algorithm S2, described in more detail in the next paragraphs). We also observed that when updating distances in the `dist` dictionary in Algorithm 1 of the main text, many of the $u \rightsquigarrow v$ pairs can remain untouched if they do not include any nodes from the newly-added `paths`$[j]$. Thus, we can calculate the distances from $u$ to the subset of target nodes $T$ for which $u \rightsquigarrow v$ needs to be recalculated, which speeds up the algorithm in practice.

`MultiTargetDijkstra()` takes four arguments: a graph $G$, a set of existing (DAG) nodes $V' \subset V$, a source node $s \in V$, and a subset of target nodes $T \subset V'$. Note that $V'$ is the set of nodes in the existing DAG (e.g., $G_0$ or $G_j$); we still need to pass in the full set of nodes from the DAG to ensure that they are not used as internal nodes on a path. The algorithm returns a distance dictionary $D_{\text{final}}$ of the distances from $s$ to each node in $T$, ensuring that a node in $T$ is never considered an internal node on a path from $s$ and returning early if all nodes in $T$ are reached.

`MultiTargetDijkstra()` differs from a standard Dijkstra algorithm in the following ways. First, we keep track of the set of nodes from $T$ that have been found (Lines 7 and 11) and we terminate the WHILE loop early if we have found all of the nodes in $T$ (Line 8). Further, we only explore dequeued node $x$'s neighbors if $x$ is not in $V'$ (e.g., if the IF statement in Line 10 is `false` and the IF statement in Line 12 is `true`). Finally, we return a dictionary $D_{\text{final}}$ containing the nodes in $T$, some of which may be infinite if they were never reached.

**Algorithm S2** `MultiTargetDijkstra`$(G = (V, E), V' \subseteq V, s \in V, T \subseteq V')$

1: $D \leftarrow \{u : \infty\}$ for all $v \in V$
2: $D[s] = 0$
3: $Q = \emptyset$
4: **for** $u \in V$ **do**
5:     Enqueue $u$ to $Q$ with priority $D[s]$
6: **end for**
7: found $\leftarrow \emptyset$
8: **while** $|T| > |\text{found}|$ and $Q \neq \emptyset$ **do**
9:     $x \leftarrow$ dequeue $Q$ with min priority
10:     **if** $x \in T$ **then**
11:         found $\leftarrow$ found $\cup\{x\}$
12:     **else if** $x \notin V'$ **then**
13:         **for** $y \in N_x$ **do**
14:             **if** $D[y] > D[x] + w_{xy}$ **then**
15:                 $D[y] = D[x] + w_{xy}$
16:                 update $Q[y]$ with priority $D[y]$
17:             **end if**
18:         **end for**
19:     **end if**
20: **end while**
21: $D_{\text{final}} \leftarrow \{v : D[v]\}$ for all $v \in T$
22: **return** $D_{\text{final}}$

### S2 Supplementary Results

#### S2.1 Topological Differences between DAG Reconstructions

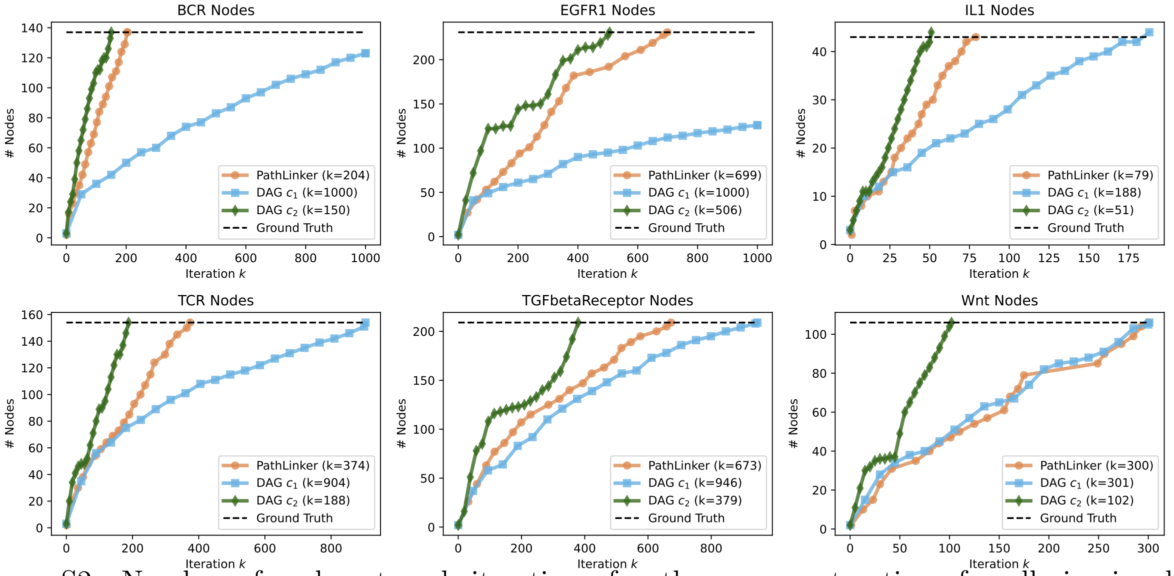

Figure S2: Number of nodes at each iteration of pathway reconstructions for all six signaling pathways. $c_1$: **min_edge_cost**; $c_2$: **min_paths_cost**.

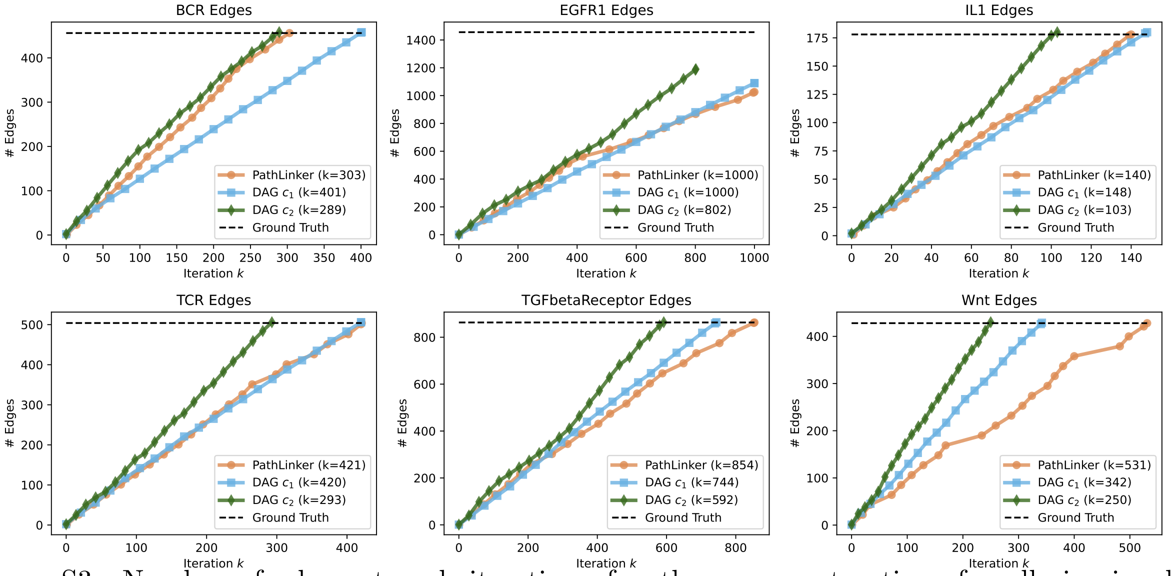

Figure S3: Number of edges at each iteration of pathway reconstructions for all six signaling pathways. $c_1$: **min_edge_cost**; $c_2$: **min_paths_cost**.

##### S2.2 Comparison to Ground Truth Pathways

| | Ground Truth | | Reconstructions | | |
|---|---|---|---|---|---|
| **Name** | Nodes | Edges | Method | Nodes ($k$) | Edges ($k$) |
| BCR | 137 | 456 | PathLinker | 137 (204) | 456 (303) |
| | | | DAG $c_1$ | *123 (1000)* | 456 (401) |
| | | | DAG $c_2$ | 137 (150) | 456 (289) |
| EGFR1 | 231 | 1456 | PathLinker | 231 (699) | *1026 (1000)* |
| | | | DAG $c_1$ | *126 (1000)* | *1089 (1000)* |
| | | | DAG $c_2$ | 231 (506) | *1189 (1000)* |
| IL1 | 43 | 178 | PathLinker | 43 (79) | 178 (140) |
| | | | DAG $c_1$ | 43 (188) | 178 (148) |
| | | | DAG $c_2$ | 44 (51) | 178 (103) |
| TCR | 154 | 504 | PathLinker | 154 (374) | 504 (421) |
| | | | DAG $c_1$ | 154 (904) | 504 (420) |
| | | | DAG $c_2$ | 154 (188) | 504 (293) |
| TGF$\beta$ | 209 | 863 | PathLinker | 209 (673) | 863 (854) |
| | | | DAG $c_1$ | 209 (946) | 863 (744) |
| | | | DAG $c_2$ | 209 (379) | 863 (592) |
| Wnt | 106 | 428 | PathLinker | 106 (300) | 428 (531) |
| | | | DAG $c_1$ | 106 (301) | 428 (342) |
| | | | DAG $c_2$ | 106 (102) | 429 (250) |

Table S1: Values of $k$ used for each method in precision and recall analysis. For each signaling pathway, the method was run until the number of ground truth nodes and/or edges were reached (the final value of $k$ required to reach this threshold is listed in parentheses). Entries in italics did not reach the ground truth number of nodes or edges when $k = 1000$.

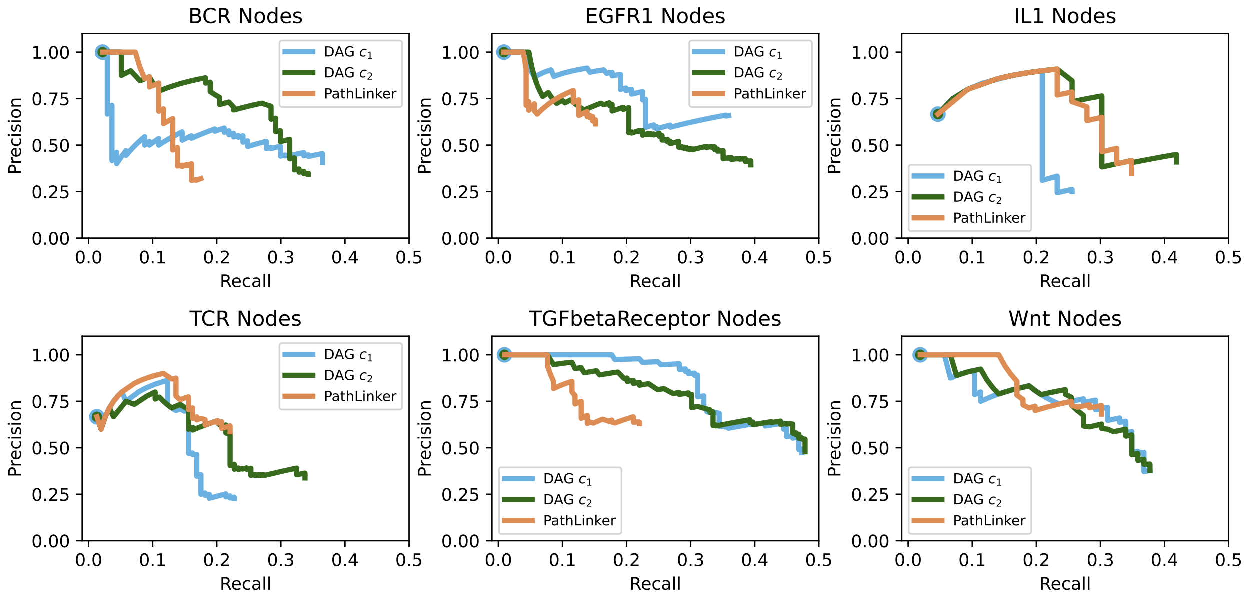

Figure S4: Precision-recall curves of nodes for size-matched reconstructions from six pathways. c1: **min_edge_cost**; c2: **min_paths_cost**.

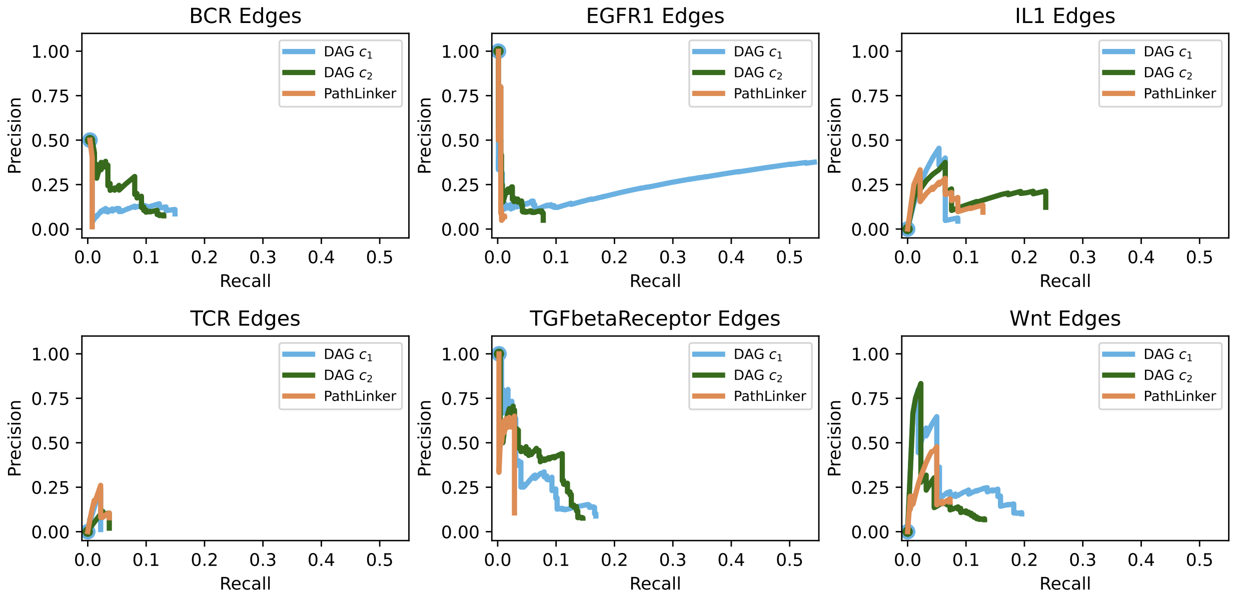

Figure S5: Precision-recall curves of edges for size-matched reconstructions from six pathways. c1: **min_edge_cost**; c2: **min_paths_cost**.

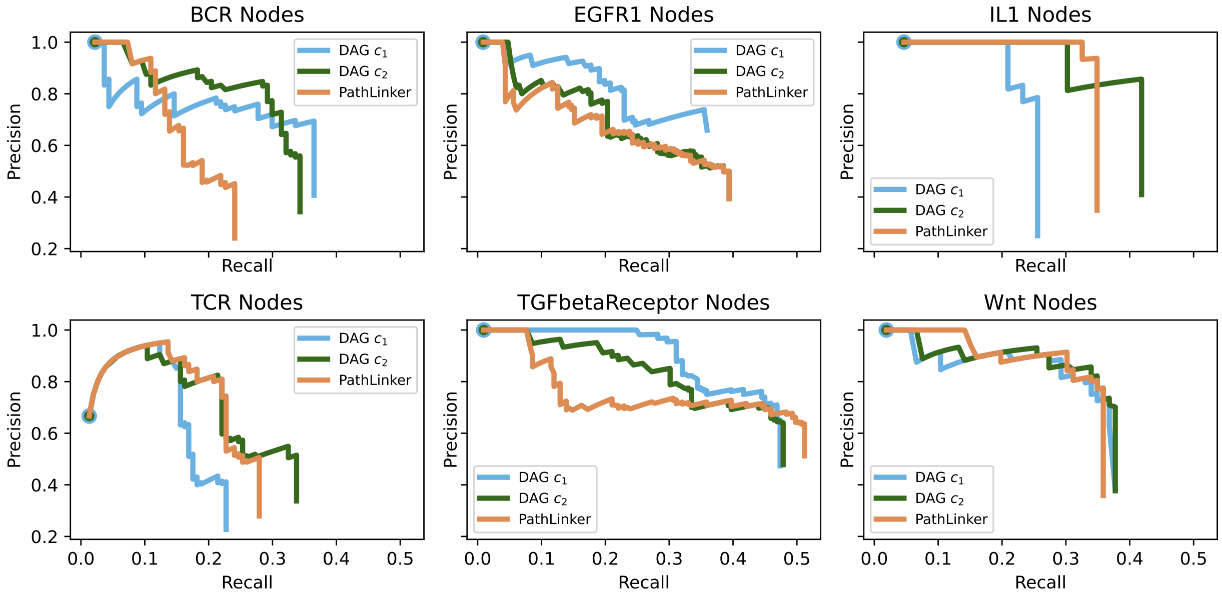

Figure S6: Precision-recall curves of nodes for reconstructions from six pathways when negatives are subsampled in a 50:1 ratio compared to positive. This is one representative sampling of ten repeated experiments. c1: **min_edge_cost**; c2: **min_paths_cost**.

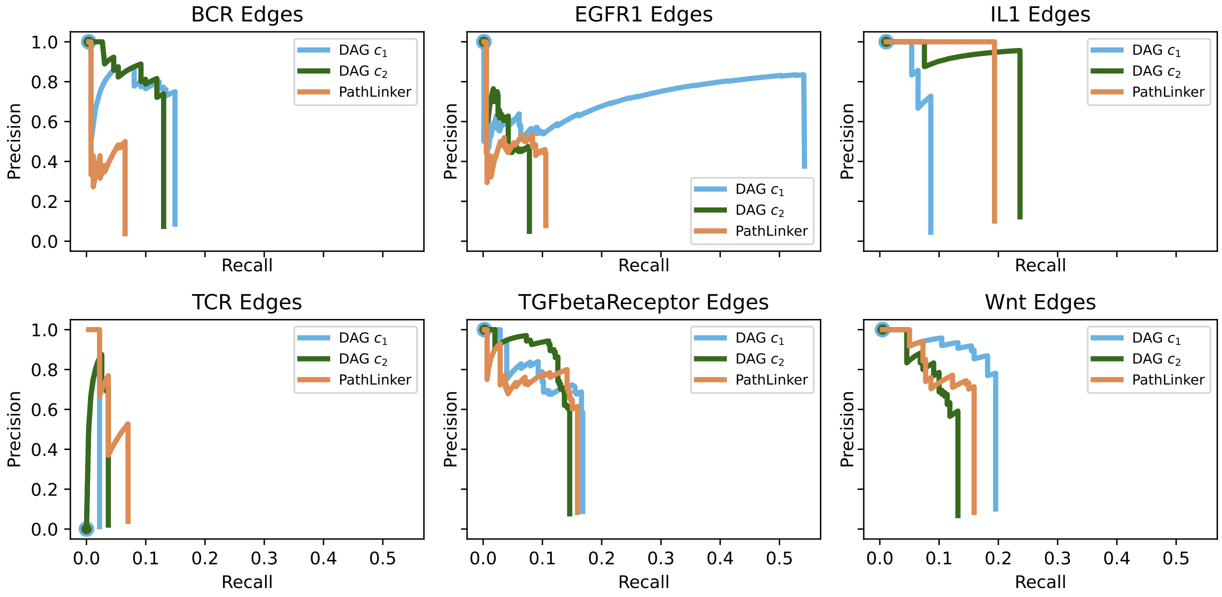

Figure S7: Precision-recall curves of edges for reconstructions from six pathways when negatives are subsampled in a 50:1 ratio compared to positive. This is one representative sampling of ten repeated experiments. c1: **min_edge_cost**; c2: **min_paths_cost**.

| **Name** | **DAG** $c_1$ | **DAG** $c_2$ | **PathLinker** |
|---|---|---|---|
| BCR nodes | $0.252 \pm 0.02$ | $\mathbf{0.281 \pm 0.01}^*$ | $0.162 \pm 0.01$ |
| BCR edges | $\mathbf{0.113 \pm 0.01}$ | $0.106 \pm 0.01$ | $0.026 \pm 0.00$ |
| EGFR1 nodes | $\mathbf{0.290 \pm 0.00}$ | $0.280 \pm 0.00^*$ | $0.267 \pm 0.00$ |
| EGFR1 edges | $\mathbf{0.378 \pm 0.01}^*$ | $0.046 \pm 0.00^*$ | $0.050 \pm 0.00^*$ |
| IL1 nodes | $0.197 \pm 0.01$ | $\mathbf{0.354 \pm 0.01}$ | $0.297 \pm 0.00$ |
| IL1 edges | $0.070 \pm 0.00$ | $\mathbf{0.212 \pm 0.01}$ | $0.162 \pm 0.02$ |
| TCR nodes | $0.159 \pm 0.01$ | $\mathbf{0.240 \pm 0.01}$ | $0.209 \pm 0.01$ |
| TCR edges | $0.015 \pm 0.00$ | $0.024 \pm 0.00$ | $\mathbf{0.042 \pm 0.00}$ |
| TGF$\beta$ nodes | $\mathbf{0.417 \pm 0.00}$ | $0.405 \pm 0.01$ | $0.396 \pm 0.01$ |
| TGF$\beta$ edges | $\mathbf{0.133 \pm 0.01}$ | $0.129 \pm 0.00$ | $0.118 \pm 0.01$ |
| Wnt nodes | $0.313 \pm 0.01$ | $\mathbf{0.325 \pm 0.01}$ | $0.312 \pm 0.01$ |
| Wnt edges | $\mathbf{0.169 \pm 0.01}$ | $0.104 \pm 0.01$ | $0.129 \pm 0.01$ |

Table S2: Mean and standard deviation of area under the precision-recall (AUPRC) values for nodes (left) and edges (right) for size-matched reconstructions when negatives are subsampled in a 50:1 ratio compared to positives (10 repeated experiments). $c_1$: **min_edge_cost**; $c_2$: **min_paths_cost**. Largest AUPRC values are shown in bold; asterisks denote reconstructions that did not reach the number of ground truth nodes or edges by $k = 1000$.

#### S2.3 Reconstructions Capture Diverse Biological Processes

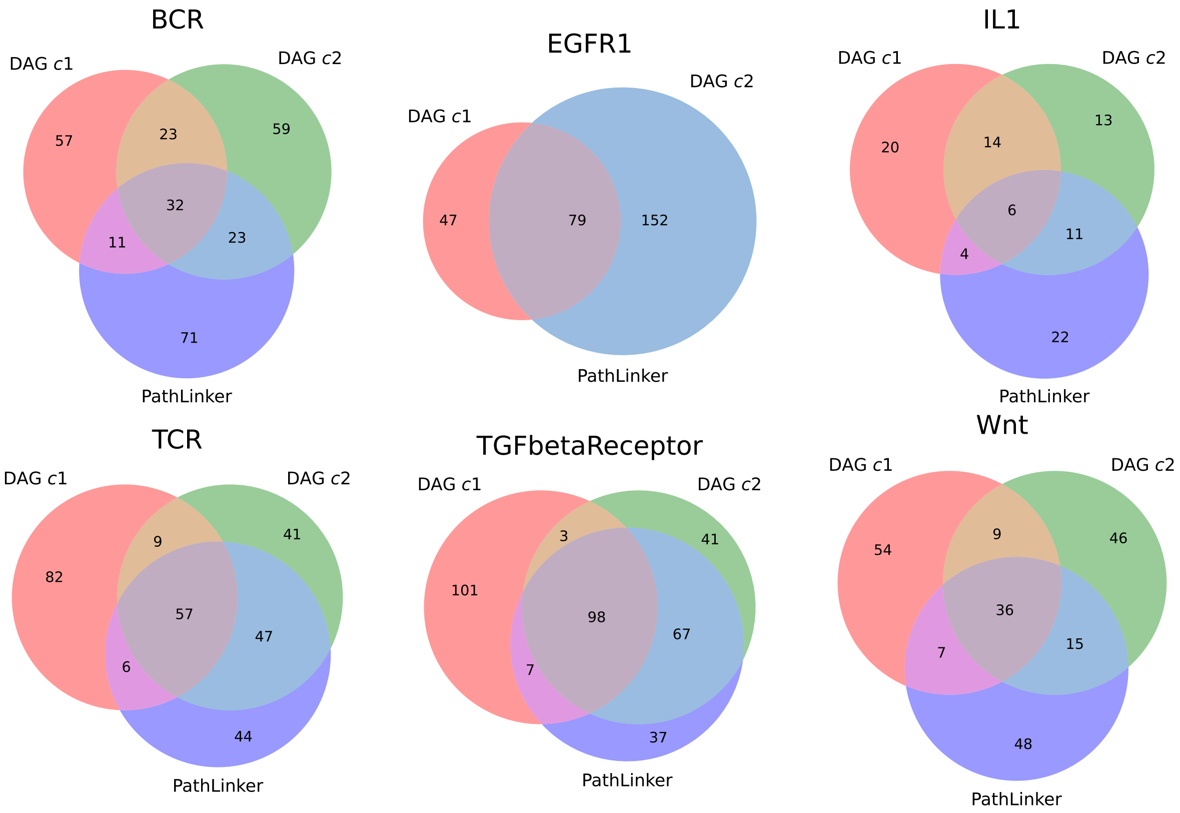

Figure S8: Venn diagram of predicted nodes for DAG **min_edge_cost**, DAG **min_paths_cost**, and PathLinker for the six pathway reconstructions with $k = 200$. c1: **min_edge_cost**; c2: **min_paths_cost**.

| **Panther Pathway** | $n$ | $k$ | **Adj. P-value** |
|---|---|---|---|
| Interferon-gamma signaling pathway (P00035) | 30 | 3 | 6.64e−04 |
| Toll receptor signaling pathway (P00054) | 61 | 4 | 2.35e−04 |
| Insulin/IGF pathway-mitogen activated protein kinase kinase/MAP kinase cascade (P00032) | 33 | 2 | 2.21e−02 |
| Parkinson disease (P00049) | 101 | 6 | 7.12e−06 |
| VEGF signaling pathway (P00056) | 68 | 4 | 3.18e−04 |
| FAS signaling pathway (P00020) | 34 | 2 | 2.22e−02 |
| B cell activation (P00010) | 71 | 4 | 3.40e−04 |
| T cell activation (P00053) | 92 | 5 | 7.79e−05 |
| Angiotensin II-stimulated signaling through G proteins and beta-arrestin (P05911) | 37 | 2 | 2.38e−02 |
| Ras Pathway (P04393) | 76 | 4 | 4.03e−04 |
| TGF-beta signaling pathway (P00052) | 102 | 5 | 7.90e−05 |
| Apoptosis signaling pathway (P00006) | 125 | 6 | 1.77e−05 |
| Gonadotropin-releasing hormone receptor pathway (P06664) | 237 | 9 | 2.52e−07 |
| Endothelin signaling pathway (P00019) | 85 | 3 | 8.86e−03 |
| PI3 kinase pathway (P00048) | 57 | 2 | 4.91e−02 |
| CCKR signaling map (P06959) | 173 | 6 | 7.34e−05 |
| Interleukin signaling pathway (P00036) | 89 | 3 | 9.54e−03 |
| Angiogenesis (P00005) | 181 | 6 | 8.10e−05 |
| EGF receptor signaling pathway (P00018) | 142 | 4 | 3.12e−03 |
| PDGF signaling pathway (P00047) | 145 | 4 | 3.17e−03 |
| Integrin signalling pathway (P00034) | 200 | 5 | 1.04e−03 |
| FGF signaling pathway (P00021) | 127 | 3 | 2.21e−02 |
| Inflammation mediated by chemokine and cytokine signaling pathway (P00031) | 261 | 5 | 3.25e−03 |
| Wnt signaling pathway (P00057) | 314 | 4 | 3.62e−02 |

Table S3: PANTHER Over-representation Test (PANTHER version 17.0, Fisher test with FDR correction of 0.05) for the nodes uniquely predicted by **min_paths_cost** for the Wnt signaling pathway reconstruction. $n$: number of proteins in the Panther pathway; $k$: number of predicted proteins in the Panther pathway.

##### S2.4 Running Times

| **Experiment** | $k$=1,...,100 | $k$=101,...,200 | **Total** |
|---|---|---|---|
| BCR $c_1$ | 0.54 | 7.3 | 7.84 |
| BCR $c_2$ | 1.17 | 22.14 | 23.3 |
| EGFR $c_1$ | 1.4 | 51.68 | 53.07 |
| EGFR $c_2$ | 0.62 | 26.13 | 26.75 |
| IL1 $c_1$ | 0.19 | 0.89 | 1.08 |
| IL1 $c_2$ | 2.53 | 24.12 | 26.66 |
| TCR $c_1$ | 0.94 | 5.94 | 6.88 |
| TCR $c_2$ | 0.6 | 2.8 | 3.4 |
| TGF$\beta$ $c_1$ | 0.2 | 0.73 | 0.93 |
| TGF$\beta$ $c_2$ | 16.26 | 44.25 | 60.51 |
| Wnt $c_1$ | 0.24 | 43.28 | 43.52 |
| Wnt $c_2$ | 0.62 | 6.14 | 6.76 |
| **Average** | 2.11 | 19.62 | 21.73 |
| **Min** | 0.19 | 0.73 | 0.93 |
| **Max** | 16.26 | 19.62 | 60.51 |

Table S4: Running times (hours) for the experiments for the first 100 iterations, the second 100 iterations, and for the total $k = 200$ iterations. Experiments were conducted on a 2020 MacBook Pro with a 2 GHz quad-core processor and 16GB memory.
